# *In situ* estimates of iron-oxidation and accretion rates for iron-oxidizing bacterial mats at Lō’ihi Seamount

**DOI:** 10.1101/095414

**Authors:** David Emerson, Jarrod J. Scott, Anna Leavitt, Emily Fleming, Craig Moyer

## Abstract

It is increasingly recognized that diffuse, hydrothermal venting is an important source of iron to the deep-sea that can influence oceanic iron dynamics and abundance. Lithotrophic Fe-oxidizing bacteria (FeOB) are dominant at diffuse hydrothermal vent sites, producing microbial iron mats that are often centimeters or more thick. At present, little is known about *in situ* Fe-oxidation rates, or accretion rates for iron mats. An *in situ* productivity chamber was developed that took advantage of the unique mineral morphotypes produced by FeOB to estimate rates of Fe-oxidation and accretion. Chambers were placed at two diffuse vents (1179 and 1300 mbsl) at Lō’ihi Seamount where they were colonized by FeOB for different amounts of time. From this analysis, it was estimated that Fe-oxidation rates could range from 8.2–51.9 × 10^−6^ mol · hr^−1^, and that iron mats could accrete at around 2.2 cm · yr^−1^. Molecular analysis indicated that the relative abundance of Zetaproteobacteria, a group of known FeOB, accounted for 80–90% of the bacteria colonizing the chambers. There was a distinct difference between populations at the 1179m site (Pohaku), and the 1300m site (North Hiolo Ridge). Microscope slides placed within the productivity chambers were colonized by different morphotypes of FeOB. The cells responsible for one common morphotype that produces a Y-shaped filament were identified as Zetaproteobacteria by use of a small subunit rRNA probe. This work confirms the importance of FeOB in the formation of chemosynthetic iron mats, and provides the first estimates for *in situ* Fe-oxidation rates and mat accretion rates.

**Highlights:**

- An *in-situ* productivity chamber was developed to estimate rates of Fe-oxidation and understand colonization patterns at chemosynthetic iron mats at Lō’ihi Seamount.
- Fe-oxidation rates ranged from 8.2–51.9 × 10^−6^ mol ˙ hr^−1^, and it was estimated that the iron mats could accrete at around 2.2 cm ˙ yr^−1^.
- The iron mat community was dominated by Zetaproteobacteria, whose relative abundance accounted for up to 89% of the microbial community.
- The community membership that grew during short-term incubations reflected the community composition of nearby microbial mats.

## 1. Introduction

Many marine hydrothermal vent systems host chemosynthetic microbial mat communities dominated by lithotrophic Fe-oxidizing bacteria (FeOB) that live on Fe(II)-rich, anoxic fluids emanating from the seafloor. These vent ecosystems are found both in the deep-sea and in shallow waters, and can be associated with active volcanic seamounts, as well as tectonic crustal spreading and subduction zones (Boothman and Lloyd, 2010; Emerson and Moyer, 2002; Kato et al., 2009; Scott et al., 2015). The most active Fe-oxidizing communities are associated with cooler, diffuse flow systems capable of supporting microbial mats that are often centimeters thick or more. Despite numerous studies of microbial iron mats, the *in situ* activities marine of FeOB are not well studied, nor do we know the accretion rates of *in situ* iron mats. The rapid heterogeneous oxidation (also referred to as auto-oxidation) of iron at circumneutral pH make it challenging to distinguish between abiotic and biotic reactions, thus making simple mass balance calculations difficult to interpret (Melton et al., 2014). Iron isotopes have proven useful for distinguishing some types of abiotic from biotic oxidation of iron, but are less suitable for specific rate measurements (Johnson et al., 2008). Thus there are inherent challenges to assessing the *in situ* activity of Fe-oxidizing microbial communities.

A more detailed understanding of the productivity of these Fe-fueled ecosystems can also help better constrain their contribution to the oceanic Fe budget. It is clear there are still significant unknowns regarding the contribution of hydrothermal sources of iron to the ocean’s iron budget (German et al., 2015; Resing et al., 2015). Furthermore, the biogenic iron oxides produced at hydrothermal vents have unique properties. They are often referred to as hydrous ferric oxides (HFO) in part because they are poorly crystalline oxides, composed primarily of ferrihydrite, with large surface areas that adsorb water and are relatively buoyant (Emerson, 2016). These biogenic oxides undergo diagenesis to crystalline forms more slowly than synthesized iron oxides, probably as a result of containing organic ligands, as well as becoming silicified (Kennedy et al., 2004; Picard et al., 2015; Toner et al., 2012). Combined, these properties of biogenic iron oxides may result in longer residence times in the water column, and increased bioavailability. There is evidence that biogenic iron oxides contribute to the iron flux into the ocean (Bennett et al., 2011; Toner et al., 2009), thus understanding details about their production is important to understanding their contribution as a global iron source.

One avenue for assessing *in situ* activity, and estimating iron oxidation rates is to take advantage of the unique morphotypes of biogenic oxides produced by FeOB. These morphotypes include cylindrical filamentous sheaths, helical stalks, short tubular stalks (referred to here as Y’s), and other filamentous forms (Chan et al., 2016). Freshly produced biogenic morphotypes are relatively easy to classify by light microscopy, thus FeOB offer a rare opportunity to observe microbial growth in nature. We chose to utilize this approach, hypothesizing that use of a chamber designed to capture iron oxides and cell morphologies could provide insight into how FeOB colonization may occur, further our understanding of the population dynamics involved in colonization, and provide an estimate for *in-situ* growth rates.

## 2. Materials and Methods

### 2.1 Chamber Design

A productivity chamber was designed for seafloor deployment. The chamber consisted of a central open rectangular section that held microscope slides and weights (Fig. 1). The microscope slides were placed in standard 50 ml conical tubes (1 slide/tube) with the bottoms removed. A zip-tie was attached at the top and bottom of the tube to prevent the slides from falling out. The conical tubes were then attached to the open interior of the chamber with zip-ties. A hollow tube, referred to here as a cassette, with open screw-cap ends that had Nitex nylon mesh cut to fit the openings on either end was strapped on either side of the chamber with zip ties, as shown in Fig.1. The volume each cassette was 95 ml. The chamber and cassette was constructed from PVC hardware purchased at a building supply store. To allow for manipulation by the robotic arm of a submersible, a lanyard of floating polypropylene line with a small float of syntactic foam was attached to each chamber.

**Fig. 1.**
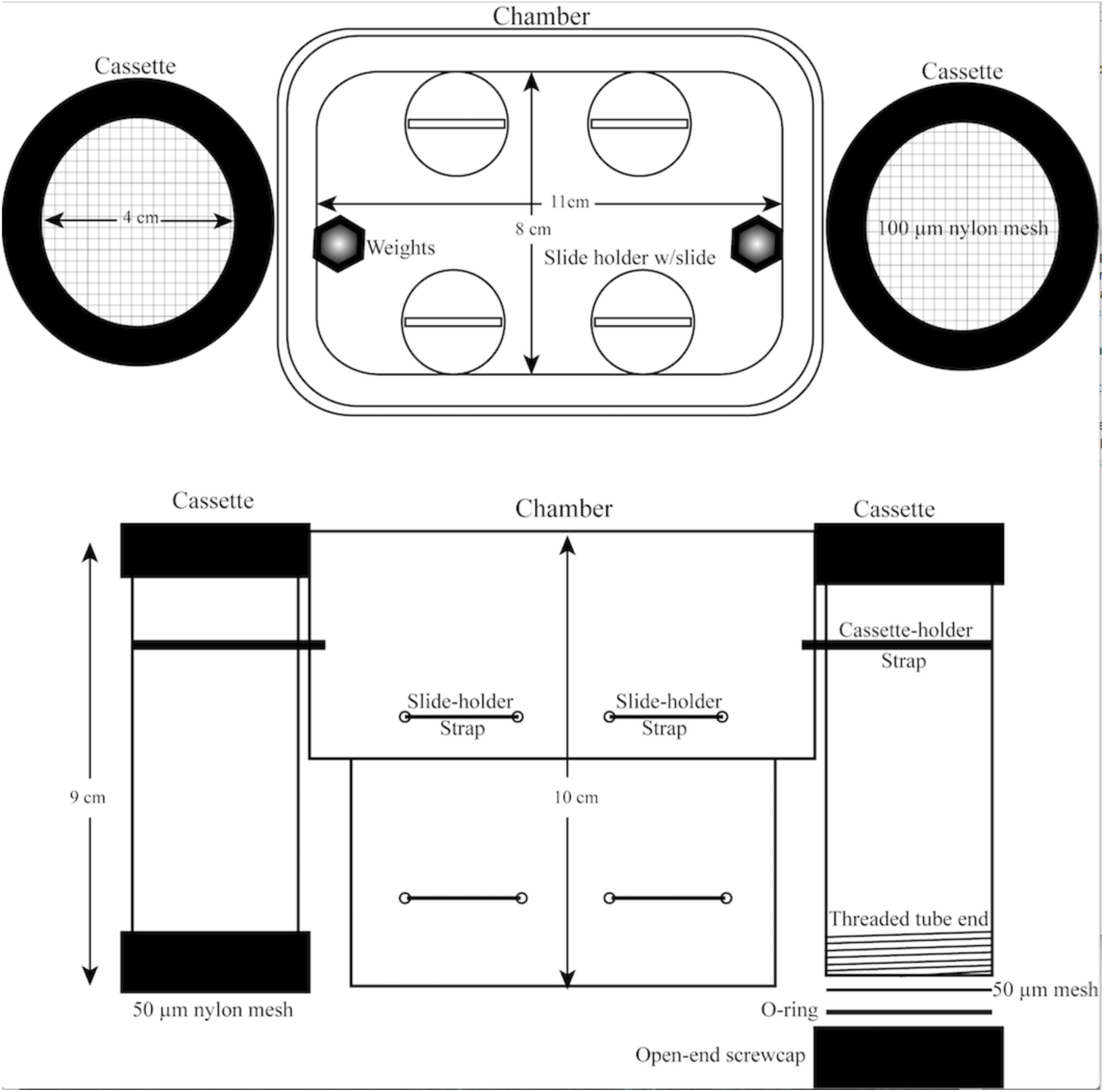
A diagram of the productivity chambers showing the central body of the chamber with that held the microscope slides, and the two cassettes, covered on both ends with Nitex mesh. The cassettes had screwcaps at both ends that were used to hold the mesh that was cut to fit the openings.

### 2.2 Chamber Deployment and recovery

Chambers were deployed at two sites on the seafloor in close vicinity to diffuse flow hydrothermal vents at Lō’ihi Seamount (18.920°N, 155.270°W). Deployment was done using the ROV *Jason 2* during the initial dives of a 14 day expedition to Lō’ihi in spring of 2013. The first deployment of 4 chambers (8 cassettes) was at Pohaku vents (sometimes referred to as Marker 57) at a depth of 1,179m. The second deployment of four chambers was at North Hiolo Ridge (NHR, Marker 39) at a depth of 1300m. Chambers were inspected on ‘fly-bys’ during subsequent submersible operations to visually assess if iron oxides were accumulating in the cassettes. At Pohaku, due to a lack of iron-oxide accumulation, the chambers were redeployed after 7 days, as described in the Results and Discussion section. Miniature temperature recorders (MTRs) were deployed in vents at both the NHR and Pohaku sites.

For recovery, the chambers were placed in a closed bio-box mounted on the submersible to maintain static water conditions during subsequent ROV operations. As soon as *Jason* was on deck, the bio-box was opened, and while still submerged rubber stoppers (size 9 1/2) were pushed into the bottom opening of the cassettes to prevent the water entrained in them from draining out after the chambers were removed from the bio-box. In the lab the cassettes were removed from the chamber body, and the top mesh removed. The iron oxides that accumulated in the cassette were placed in sterile 50 ml conical tubes with a 25 ml pipette. These oxides were processed for analysis of cell counts, morphology, and total iron, and a subsample was frozen for DNA extraction. The microscope slides from the chamber interior were removed and air-dried. Perhaps as a result of the longer incubation times at NHR, the individual cassettes for each chamber contained more oxides than the Pohaku chambers. For this reason, each cassette from the NHR chambers was harvested separately, while the two cassettes for each of the four Pohaku samples were combined to a single sample. As a result, the oxidation rates reported in Table 1 are broken into the individual cassettes for NHR samples, but combined for the cassettes from the Pohaku chambers.

**Table 1.** Fe-oxidation rates at NHR and Pohaku. The values for chambers for NHR were calculated from each individual cassette, while values for Pohaku are from the combined cassettes from each chamber. Chamber 6 from NHR is not included in this analysis.

| Chamber | Site | Hours | Total Fe<br>(10<sup>-6</sup> mol) | Total cells | Fe-oxid<br>10<sup>-6</sup> mol · hr<sup>-1</sup> | Fe-oxidized per cell<br>10<sup>-16</sup> mol · hr<sup>-1</sup> · cell<sup>-1</sup> |
| --- | --- | --- | --- | --- | --- | --- |
| ST1a | NHR | 148 | 3882 | 1.70E+08 | 26.2 | 1.54 |
| ST1b | NHR | 148 | 7674 | 3.59E+08 | 51.9 | 1.44 |
| ST7a | NHR | 148 | 6229 | 4.17E+08 | 42.1 | 1.01 |
| ST7b | NHR | 148 | 7125 | 4.33E+08 | 48.1 | 1.11 |
| **Average (St Dev)** |  |  | **6228 (1673)** | **3.5E+08 (1.1E+08)** | **42.1 (11.3)** | **1.3 (.26)** |
| ST2a | NHR | 256 | 3966 | 4.14E+08 | 15.5 | 0.38 |
| ST2b | NHR | 256 | 2090 | 7.97E+07 | 8.2 | 1.05 |
| **Average (St Dev)** |  |  | **3028 (1326)** | **2.5E+08 (2.4E+08)** | **11.8 (5.2)** | **0.72 (0.4)** |
| ST3 | Pohaku | 112 | 919 | 5.05E+07 | 8.2 | 1.63 |
| ST4 | Pohaku | 112 | 2070 | 8.74E+07 | 18.5 | 2.04 |
| ST5 | Pohaku | 112 | 2082 | 7.72E+07 | 18.6 | 2.31 |
| ST8 | Pohaku | 112 | 1686 | 1.24E+08 | 15.1 | 1.12 |
| **Average (St Dev)** |  |  | **1689 (546)** | **8.5E+07 (3.1E+07)** | **15.1 (4.9)** | **1.8 (0.53)** |

### 2.3 Cell counts

Cell counts were done on aliquots of the iron oxides that were fixed in 2% glutaraldehyde. Counting was done by diluting the original sample between 1:2 and 1:5, depending upon the initial cell density, and then using 10 μl of the diluted material to make a uniform smear within a 1 cm diameter ring on an agarose-coated microscope slide. This was allowed to air-dry, and then 6 μl of d-H_2_O plus 3 μl of the nucleic acid dye Syto13 (Invitrogen) was added, and the cells were counted by epifluorescence microscopy using the 100x objective on an Olympus BX60 microscope. Further details on these procedures can be found in (Emerson and Moyer, 2002).

### 2.4 Morphological analysis

To assess the different morphotypes of iron oxides present in the cassettes, the cell counting samples were subjected to an additional 1:10 dilution and prepared on the counting slides, as described above. Additional dilution ensured minimal overlapping of individual oxide particles on the microscope slide. Images of 8 random microscope fields (40x objective, brightfield) were collected for three different aliquots of the same sample. Imaging was done using a Qicam Fast CCD camera (QImaging, Surrey, BC, Canada) mounted on an Olympus BX60 microscope. The image analysis program ImageJ (https://imagej.nih.gov/nih-image/) was used to quantify the different morphotypes in each image. Morphotypes were classified as sheaths, stalks, Y-shaped structures, filaments, and amorphous oxide particles of no determinate shape. Details on the image analysis and definition of morphotypes has been published elsewhere (Scott et al., 2015). Analysis of colonization slides was done by light microscopy. To qualitatively assess coverage and the different morphotypes present on the slides, arbitrary images (40x objective) were taken at 0.5 cm intervals for the entire length of the 7.5 cm microscope slide.

To make representational images of different individual morphotypes of either stalks, Y’s, or stick-like filaments, incubation slides were stained with Syto13 and examples of different morphotypes were imaged, either as individual photomicrographs, or several overlapping images were used to make a photomosaic in ImageJ. These images were used to determine the length of individual filaments, either stalks or Y’s as reported below, as well as create composite fluorescent and light images to show the juxtaposition of cells producing Fe-oxide structures.

### 2.5 Total iron

The total amount of Fe-oxides that had accumulated in each cassette, or set of 2 cassettes was determined by diluting a sample of the oxides 1:10 in 0.1 M hydroxylamine to reduce the oxides to Fe(II). This solution was then diluted 1:10 or 1:20 into a ferrozine solution to colorimetrically quantify the amount of iron present. Iron was assayed using the ferrozine method (Stookey, 1970) (34) adapted for reading on a Mikroscan 96 well plate reader (Mcbeth et al., 2011).

### 2.6 Fluorescent in situ Hybridization (FISH)

Phylogenetic staining was used to identify the phylogeny of cells attached to Y-shaped iron-oxides. Samples enriched in this morphotype were collected, preserved with paraformaldhyde, washed with PBS, and resuspended in 1:1 PBS ethanol at -20 °C until further analysis. Cells attached to the terminal end of Y shaped iron oxides was performed using Catalyzed Activated Reporter Deposition FISH (CARD-FISH; (Pernthaler et al., 2002)) and modified to limit the non-specific binding of probes that occurs in iron-oxide rich samples with several blocking agents (Fleming et al., 2013). Finally, a Cy-3 Zeta proteobacteria specific probe, unlabeled helper probes, and a 20% formamide concentration (Fleming et al., 2013) were used to confirm the phylogenetic affiliation of the Y attached cells.

### 2.7 Community analysis

We selected 6 samples for 454-pyrosequencing analyses, two from NHR–148h (chambers 1 & 7), two from NHR–256h (chambers 2 & 6), and two from Pohaku (chambers 5 & 8). Samples were processed following previously published protocols (Scott *et al.*, 2015). From each mat sample, approximately 250 mg (wet weight) of mat material was used for DNA extraction from each sample using a Mo Bio PowerSoil^®^ DNA Extraction Kit (Mo Bio Laboratories, Carlsbad CA, USA), modified to include an initial phenol:chloroform:isoamyl alcohol (PCI) step. We found that adding an initial PCI treatment increased final DNA yield from an average < 5ng/μL to concentrations typically > 15 ng/μL (data not shown). Briefly, 200 μL of bead solution was removed from each bead tube and replaced with 200 μL of 25:24:1 PCI (Sigma-Aldrich, St. Louis MO, USA). Samples were then extracted using the manufacture recommended protocol and sent to Research and Testing Laboratory (Lubbock TX, USA) for pyrosequencing. We targeted the V4–V5 hypervariable region of the small subunit rRNA gene (SSU rRNA; *E. coli* positions 531–997) using 530F (5'-GTG CCA GC**M** GC**N** GCG G-3') and 1100R (5'-GGG TT**N** CG**N** TCG TTG-3') following established protocols (Dowd et al., 2008).

Sequence processing was performed using mothur v.1.35.0 (Schloss et al., 2009) following previously published methodology (Schloss et al., 2011). Specifically, we used mothur to remove primer and barcode sequences from all reads, as well as any short reads (< 250 bp), reads containing more than six homopolymers, reads with any ambiguities, and/or any chimeric reads detected by UCHIME (Edgar et al., 2011). All quality filtered reads were aligned against a mothur-compatible re-creation (mothur.org/wiki/Silva_reference_files; last accessed 09.11.2016) of the SILVA-SEED (SILVA v123) reference alignment (Quast et al., 2012). Pyrotag reads were classified against the Greengenes reference taxonomy (McDonald et al., 2012). All pyrosequencing libraries are deposited at the European Nucleotide Archive under the sample accessions numbers ERS1466444–ERS1466449, study accession number PRJEB18515. For comparative purposes we included data from iron mats at the Mid-Atlantic Ridge and Lō’ihi (Scott et al., 2016).

### 2.8 Community analysis

Minimum Entropy Decomposition (MED) (Eren et al., 2014) analysis was performed on the complete dataset of 142,025 reads from 52 samples using the following screening parameters: minimum substantive abundance, 28; minimum entropy value for decomposition, 0.0965; maximum number of discriminants to use for decomposition, 4; maximum variation allowed in each node, 3 nucleotides. Per node normalization was conducted before decomposition based on the node size of the most abundant sequence in the dataset (Eren et al., 2014). Screening removed 18,290 outliers resulting in 123,735 total reads in the final analysis. Alignments contained 570 characters with an average read length of 268 base pairs without gaps. Bray-Curtis dissimilarity coefficient (Bray and Curtis, 1957) was used to calculate distance metrics for non-metric multidimensional scaling (NMDS) analyses.

In addition, all reads from MED nodes classified as Zetaproteobacteria were classified further using ZetaHunter (https://github.com/mooreryan/ZetaHunter, last accessed 12.05.2016)—a command line script designed to assign SSU rRNA gene sequences to Zetaproteobacteria OTUs (97% identity) (Edgar et al., 2011; Schloss et al., 2009) defined by a reference database with an underlying phylogenetic structure (SILVA v123) (Quast et al., 2012; Pruesse et al., 2012). In the absence of a well-defined taxonomy, OTU binning is currently the accepted method for classifying diversity of Zetaproteobacteria (McAllister et al., 2011), henceforth referred to as ZetaOtus.

## 3. Results and Discussion

### 3.1 Chamber colonization

The basic design premise for the productivity chambers was that Fe(II)-rich fluids would pass into, and through the fine mesh that enclosed both ends of the cassettes. A portion of the Fe(II) is oxidized inside the cassette resulting in precipitation of Fe-oxides that are large enough (> 100 μm) to be retained by the mesh, thus providing an estimate of accumulation rates for these oxides. A slightly larger mesh opening (100 μm) was used on the cassette exit than on the entrance (50 μm), to reduce the likelihood of back-pressure that could impede flow through the cassette. All the chambers accumulated Fe-oxides; however placement of the chambers on the topologically uneven seafloor, and in the diffuse flow venting proved challenging. At NHR (Mkr 39), four chambers were deployed around the same vent site (Supplemental Fig. 1); two were collected after 148h (6d) and the remaining two were collected at 256h (10.5d). At some point between deployment and 148h, one chamber (#6) fell away from the vent where it was placed, and was redeployed at 148h. Because we do not know the time it was out of the vent flow, this chamber was not used for rate measurements (see below). The four chambers placed at Pokahu (Mkr 57) were inspected after 7d on the seafloor, and there was no visual evidence of Fe-oxides having formed in the cassettes. Therefore, the chambers were re-deployed to another area of flow about 2 m away, and collected after 112h (4.5d). During the intervening period, visible iron oxidation had occurred in all the cassettes associated with these four chambers. The temperature at the vent associated with the Pohaku samplers was constant at around 22°C during the deployment period (Supplemental Fig. 2). At NHR, the temperature was constant for 4 d at 36.5 °C, but then decreased to 25°C and showed some fluctuation between 23.5 and 24° until recovery at 6 d (144h), indicating the MTR may have become partially dislodged from the vent (Supplemental Fig. 2). The Fe(II) and O_2_ concentrations in the chambers themselves were not measured. The Fe(II) concentrations at NHR vents are between 350 and 450 μM, while at Pohaku the Fe(II) concentrations range from around 400 to >600 μM (Glazer and Rouxel, 2009; Scott et al., 2016). These values were measured directly in vent orifices. Due to mixing with seawater there is a rapid drop in Fe(II) concentration away from the vent orifice, thus it would be expected that Fe(II) concentrations in the cassettes could be 5 – 10-fold less than orifice concentrations. Because the summit of Lō’ihi is in an oxygen minima zone, the ambient O_2_ concentrations are substantially less than fully oxygenated seawater. In 2013, the O_2_ concentration around NHR was 35 – 38 μM, while at Pohaku it was around 58 μM (Scott et al., 2016).

We currently recognize three morphotypes of biogenic Fe-oxides that are associated with specific groups of FeOB (Chan et al., 2016). These are helical stalks of the type formed by *Mariprofundus ferrooxydans*, long, round, tubular sheaths produced by filamentous cells belonging to an as yet uncharacterized Zetaproteobacteria, and flattened tubular structures that branch in a characteristic Y-shape, and are referred to here as Y’s. The sampling process often breaks these different filamentous oxides into smaller pieces. This, combined with continued mineralization through auto-oxidation of Fe(II) (e.g. (Rentz et al., 2007)), can make it difficult to distinguish individual filaments as a stalk, Y, or sheath, therefore a general filamentous oxide category is also included. Particulate oxides with no definitive morphology can also be common; however, because these resemble abiotically formed oxides, by themselves, they cannot be used as a proxy for biogenic Fe-oxide production. The percentages of the different morphotypes found in the productivity cassettes are shown in Fig. 2. This analysis revealed that recognizable biogenic morphotypes: stalks, Y’s, or filaments accounted for the majority (50 – 70%) of oxides at NHR. The exception was chamber 6 in which biogenic and particulate oxides were nearly equal. As noted above, this chamber was temporarily out of the vent flow, which may have led to less biological oxidation (it also had the least abundance of Zetaproteobacteria, see below). By contrast, the Pohaku chambers were dominated by particulate Fe-oxides, and undefined filamentous oxides accounted for most of the remainder.

**Fig. 2.**
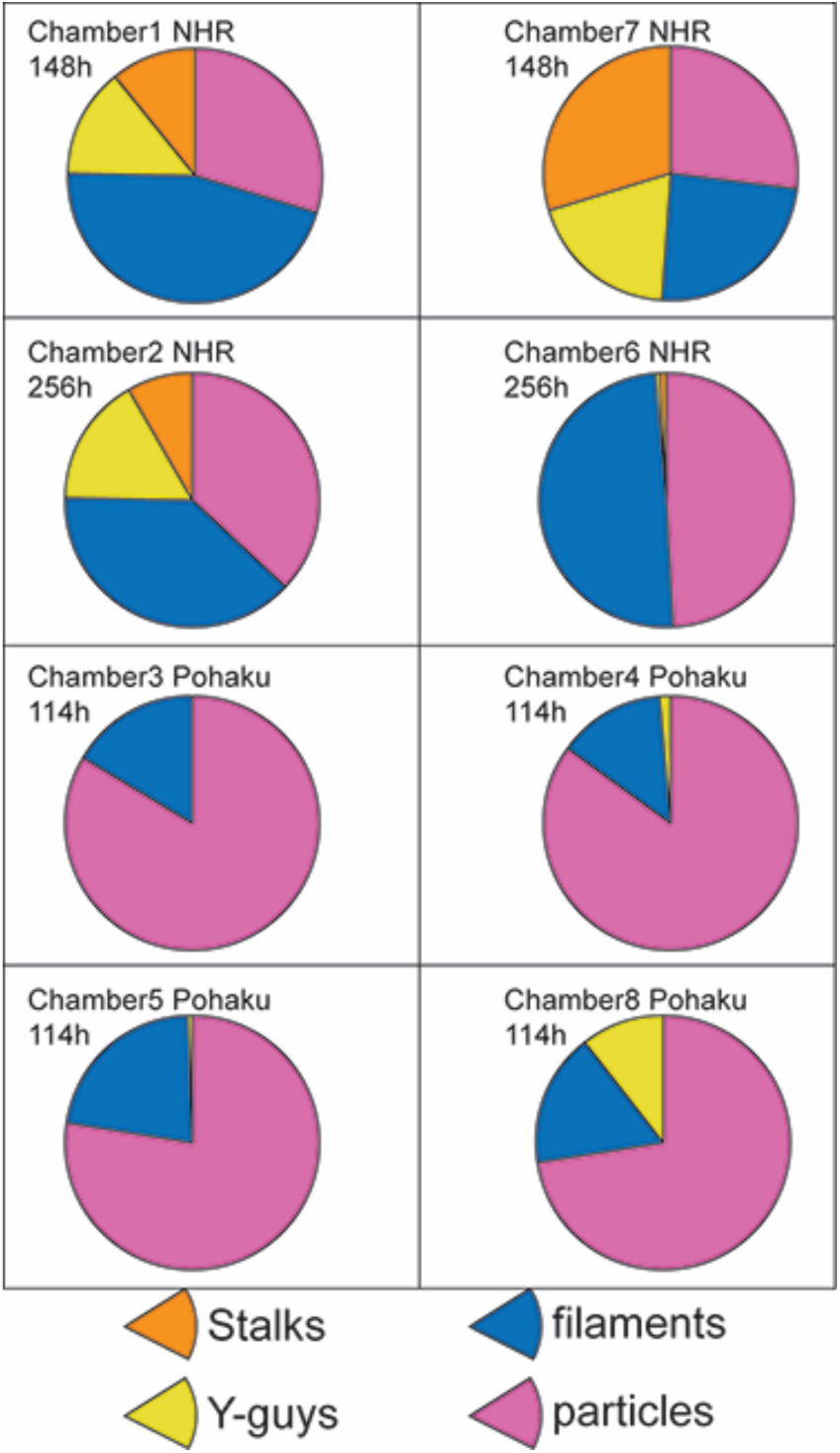
Pie charts showing the percentages of different morphotypes of iron oxides from the different cassettes. The oxide morphology was determined by light microscopy as described in the text.

### 3.2 Iron oxidation rates

Collecting the iron oxides that accumulated in each cassette revealed the short term incubations at NHR had the greatest accumulation of total iron, an average of 6.2 × 10^−3^ mol Fe, compared to an average of 3.1 × 10^−3^ mol for the long term incubation. The shorter term (114h) Pohaku samplers accumulated an average of 1.7 × 10^−3^ mol Fe, Table 1. From these values an iron oxidation rate expressed in mol · hr^−1^ was calculated:

Fe-ox rate = Total Fe Chamber (moles)/Deployment time (hr)

The results for each chamber are shown in Table 1; the rates ranged from a lower value of 8.2 × 10^−6^ mol · hr^−1^ (Pohaku) up to 51.9 × 10^−6^ mol · hr^−1^ (NHR/148hr). The Fe-oxidation rates measured at NHR were 3-fold faster for the 148h deployment than for the 256h deployment: 42.1 × 10^−6^ mol · hr^−1^ vs. 12.9 × 10^−6^ mol. hr^−1^, and the average Pohaku rate, 15.1 × 10^−6^ mol · hr^−1^ was similar to the single longer term NHR deployment (chamber #2).

The total number of cells associated with the Fe-oxides in the cassettes ranged from 5 × 10^7^ to 4.3 × 10^8^, Table 1. Because the 148h incubation at NHR was dominated by recognizable biogenic oxides, and had the most rapid accumulation of oxides, it serves as the best proxy for estimating an Fe-oxidation rate on a per cell basis. The total amount of iron accumulated in these cassettes divided by the total cell number yields an average value of 1.3 × 10^−16^ mol · cell^−1^ · hr^−1^ (SD = .26 × 10^−16^ mol · cell^−1^ · hr^−1^). This assumes that all the cells counted were FeOB, which is almost certainly not the case. Our amplicon analysis (see below) suggested the relative abundance of Zetaproteobacteria in these chambers was around 80%. If we assume this is a reasonable estimate, and that all the Zetaproteobacteria were oxidizing Fe(II) this gives a value of 1.6 × 10^−16^ mol · cell^−1^ · hr^−1^. An earlier analysis of Fe-oxidation by a pure culture of *M. ferrooxydans* estimated that cell growth required 9.16 × 10^−15^ mol Fe(II) per cell (Chan et al., 2010). This assumed a 12h doubling time for *M. ferroxydans*, and at this rate this would equate to an Fe-oxidation rate of 7.7 × 10^−16^ mol · cell^−1^ · hr^−1^. An estimate for the rate of Fe-oxidation for *M. ferrooxydans* grown on a cathode, where uptake of electrons was used as a proxy for Fe-oxidation gave a significantly higher rate of 7.5 × 10^−14^ mol · cell^−1^ · hr^−1^ (Summers et al., 2012). The *in-situ* cell-based Fe-oxidation rates from the cassettes is also less than estimates of Fe-oxidation rates for pure cultures of freshwater FeOB that range from 0.8 − 14 x 10^−14^ mol · cell^−1^ · hr^−1^ (Chan et al, 2016). It is not surprising the *in situ* rates reported here are less than laboratory rates, since the latter represent the more ideal conditions of controlled growth. Because the *in situ* growth chambers are designed as flow through devices, presumably there was a net loss of cells and Fe-oxides from the cassettes during the incubation period that would result in a reduced rate of oxidation. On the other hand, a lower per cell rate of Fe-oxidation could also suggest FeOB in these natural populations may be more efficient at oxidizing Fe(II) than the isolates.

### 3.3 Surface Colonization

All the colonization slides placed within the chambers at both NHR and Pohaku showed evidence for formation of biogenic oxides (results not shown). The most common biogenic morphotypes were Y’s and helical stalks. Sheath morphotypes were rare; nor were sheaths observed in the cassettes, indicating sheath-forming Zetaproteobacteria did not colonize the chambers. At Loihi, sheath-forming Zetaproteobacteria are responsible for forming mats with a veil-like appearance. These veil-like mats form farther from vent fluid sources (e.g. in cooler regions with more O_2_ and less Fe(II) compared to mats where stalks and Y’s predominate (Scott et al., 2016).

The cells that produce the Y-structures are easily detached during sampling (Chan et al., 2016), and the freshly produced Y-type structures that were plentiful in the cassettes had few apical cells associated with them. However, on the colonization slides the apical cells responsible for tube formation were relatively common, Fig. 3. Presumably direct attachment of the structures to the slides resulted in less disturbance and loss of cells upon sample recovery. The total length of individual Y structures on colonization slides ranged from 5.2 – 49.5 μm with a median length of 11.2 μm (SD = 2.6), Table 2. We estimated the rate of Y-filament formation by selecting the longest filaments that were observed on a given set of slides and dividing by the deployment time. The most rapid production rate for Y’s was 0.34 μm · hr^−1^ (Pohaku), while the average was 0.19 μm · hr^−1^ (SD = 0.11 μm · hr^−1^). To demonstrate that these presumptive Fe-oxidizers were members of the Zetaproteobacteria, a sample from NHR with a good representation of cells coupled to filaments was stained with a FISH probe specific for the Zetaproteobacteria. This resulted in staining of the terminal cells, thus confirming these cells are Zetaproteobacteria, Fig. 4.

**Fig. 3.**
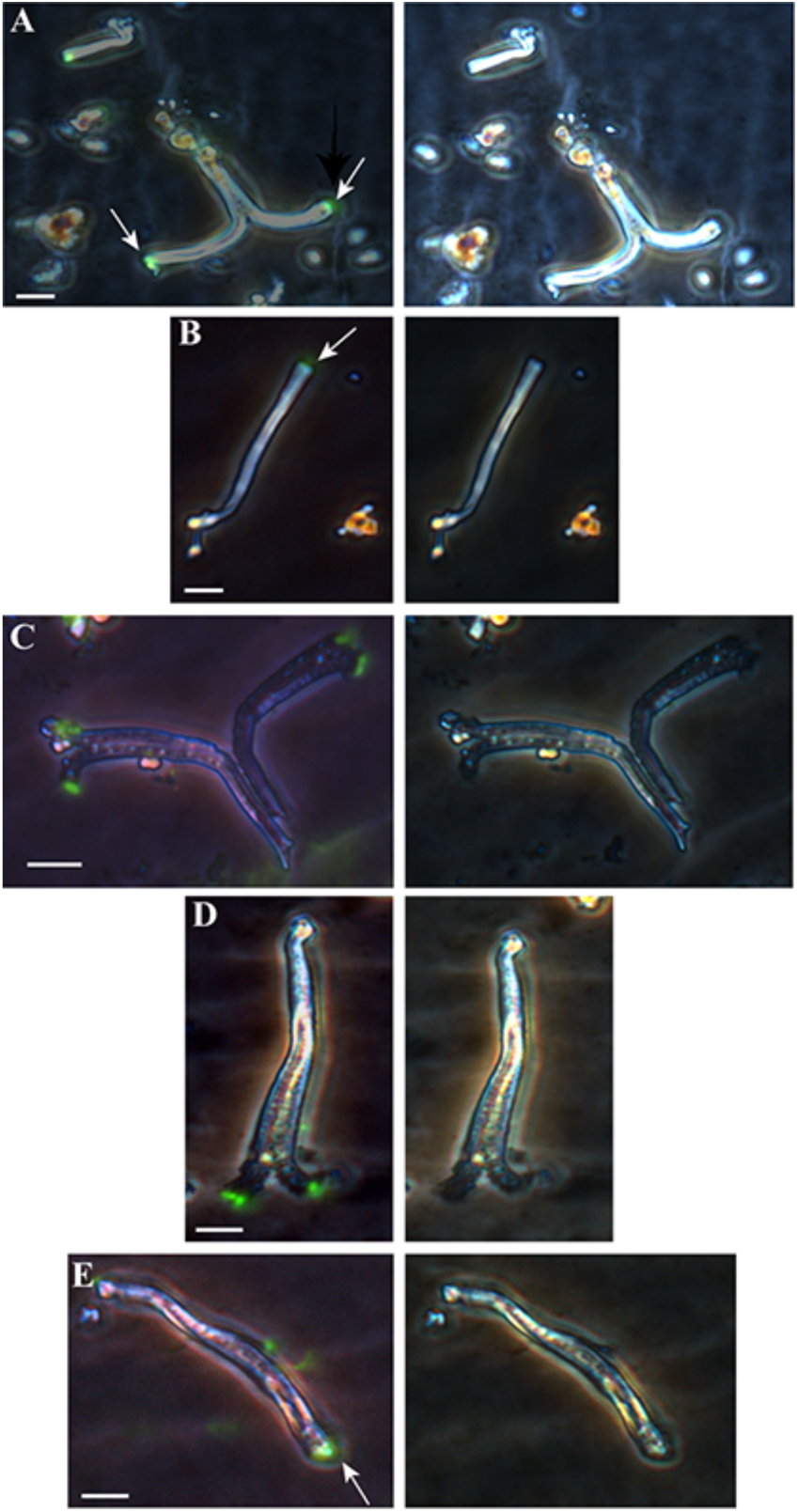
A photo-montage from an *in-situ* microscope slide of Y-shaped filaments with cells associated with the ends of the filaments. The left set of panels are overlays of epifluorescence and phase contrast images, cells are stained green at the end of the filaments, denoted by arrows in a, b, and e. The right hand panel are the original phase contrast images, note cells are not visible. The marker bar = 5 *μ*m.

**Table 2.** Filamentous iron oxide production rates for stalks and Y’s, calculated from measurements made on colonization slides.

| **Stalks** |  |  |  |  |  |  |  |  |  |
| --- | --- | --- | --- | --- | --- | --- | --- | --- | --- |
| Chamber | Site | Hrs | n | Filament length ($\mu\text{m}$) | Median length ($\mu\text{m}$) | SD | Average length top 10% | Rate $\mu\text{m/hr}$ | SD |
| 1 | NHR | 148 | 16 | 76 - 390 | 148 | 77 | 307 | 2.1 |  |
| 7 | NHR | 148 | 22 | 25-277 | 83 | 46 | 207 | 1.4 |  |
| 6 | NHR | 256 | 19 | 49-162 | 95 | 39 | 177 | 0.7 |  |
| 2 | NHR | 256 | 20 | 45-200 | 89 | 40 | 171 | 0.7 |  |
| 3 | Pohaku | 112 | 18 | 58-207 | 115 | 43 | 191 | 1.7 |  |
| **Average** |  |  |  |  | **106** | **26** |  | **1.32** | **0.62** |
| **Y's** |  |  |  |  |  |  |  |  |  |
| 1 | NHR | 148 | 28 | 7.3-15.5 | 11.9 | 2.5 | 15 | 0.1 |  |
| 2 | NHR | 256 | 19 | 5.2-17.3 | 7.8 | 3.4 | 16 | 0.06 |  |
| 7 | NHR | 148 | 16 | 6.5-15.7 | 9.5 | 2.4 | 15 | 0.1 |  |
| 5 | Pohaku | 112 | 24 | 9.0-27.0 | 14 | 5.4 | 28.3 | 0.25 |  |
| 3 | Pohaku | 112 | 27 | 8.0-49.5 | 14.2 | 8.3 | 38.3 | 0.34 |  |
| 8 | Pohaku | 112 | 11 | 9.0-27 | 9.7 | 6.4 | 29 | 0.26 |  |
| **Average** |  |  |  |  | **11.2** | **2.6** |  | **0.185** | **0.11** |

**Fig. 4.**
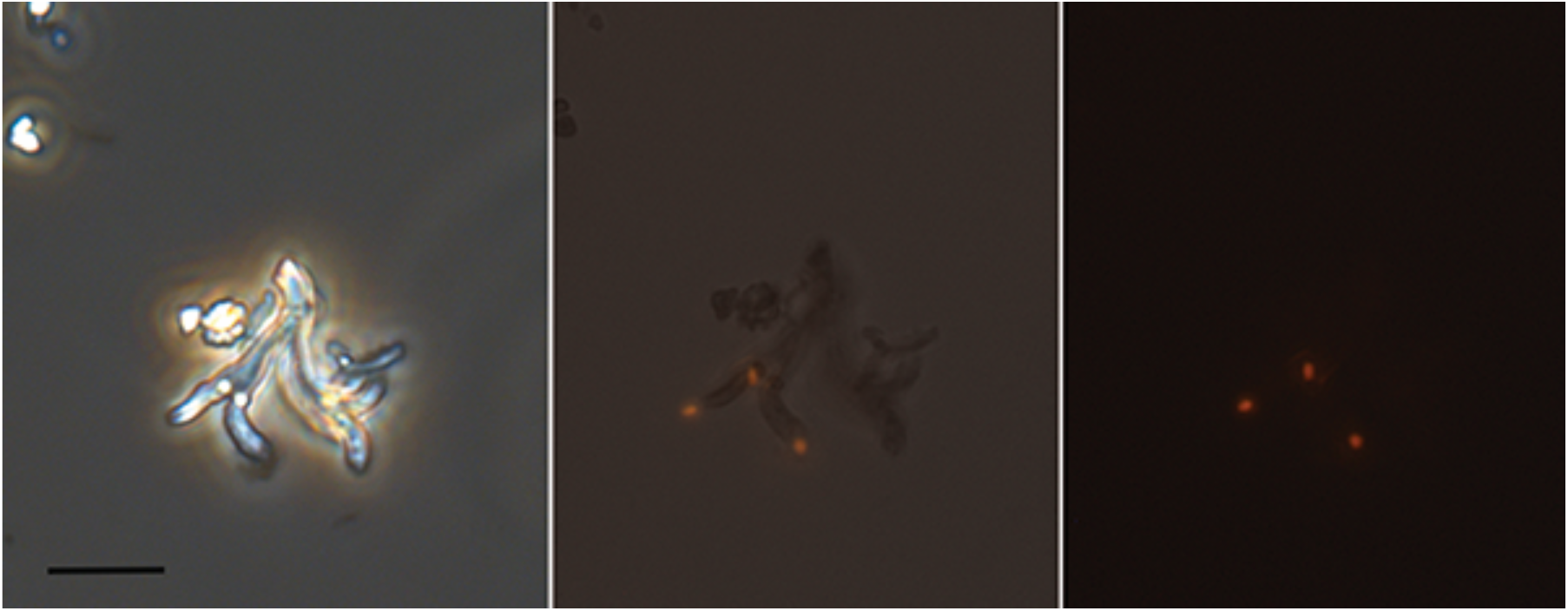
FISH-probe for Zetaproteobacteria staining apical cells in Y-shaped structures. Panel a, phase contrast image of the Y-shaped structure, the cells are not visible; b, composite epifluorescence and brightfield image of the particle; c, the epifluorescence image of FISH-probe showing cells alone. The scale bar = 5 μm.

Stalks, similar to those produced by *M. ferrooxydans*, were also observed on the colonization slides, Supplemental Fig. 3, and single stalks that had at least one cell division were common. The longest stalks could be 100’s of μm in length, with a range measured between 25 – 390 μm for individual stalks. Using the same criteria as described for the Y’s above, stalk production rates were estimated to range from 0.7 – 2.1 μm · hr^−1^, with an average of 1.32 μm · hr^−1^ (SD = 0.62 μm · hr^−1^), table 2.

A novel cell/oxide morphotype, with a stick-like appearance, was also documented on the colonization slides. This was a very thin rod <0.5 μm in diameter coated in iron oxyhydroxides, see Fig. 5. These stick-like iron oxide structures are commonly observed in marine iron mats (D. Emerson, unpublished observations), and in our morphological analysis are grouped with filamentous oxides of unknown provenance. So far as we are aware, this is the first time that cells have been observed associated with these structures in a way that indicates their genesis, and suggests this could be a novel group of FeOB. Alternatively, they could be cells that are simply being encased in Fe-oxyhydroxides as a result of abiotic Fe-oxidation. The majority of the stick-like structures did not have cells present; however this is consistent with other biogenic oxides where the majority of the structures are uninhabited (Chan et al., 2016). We were unable to visualize any of these cells using FISH, so we do not know if they belong to the Zetaproteobacteria.

**Fig. 5.**
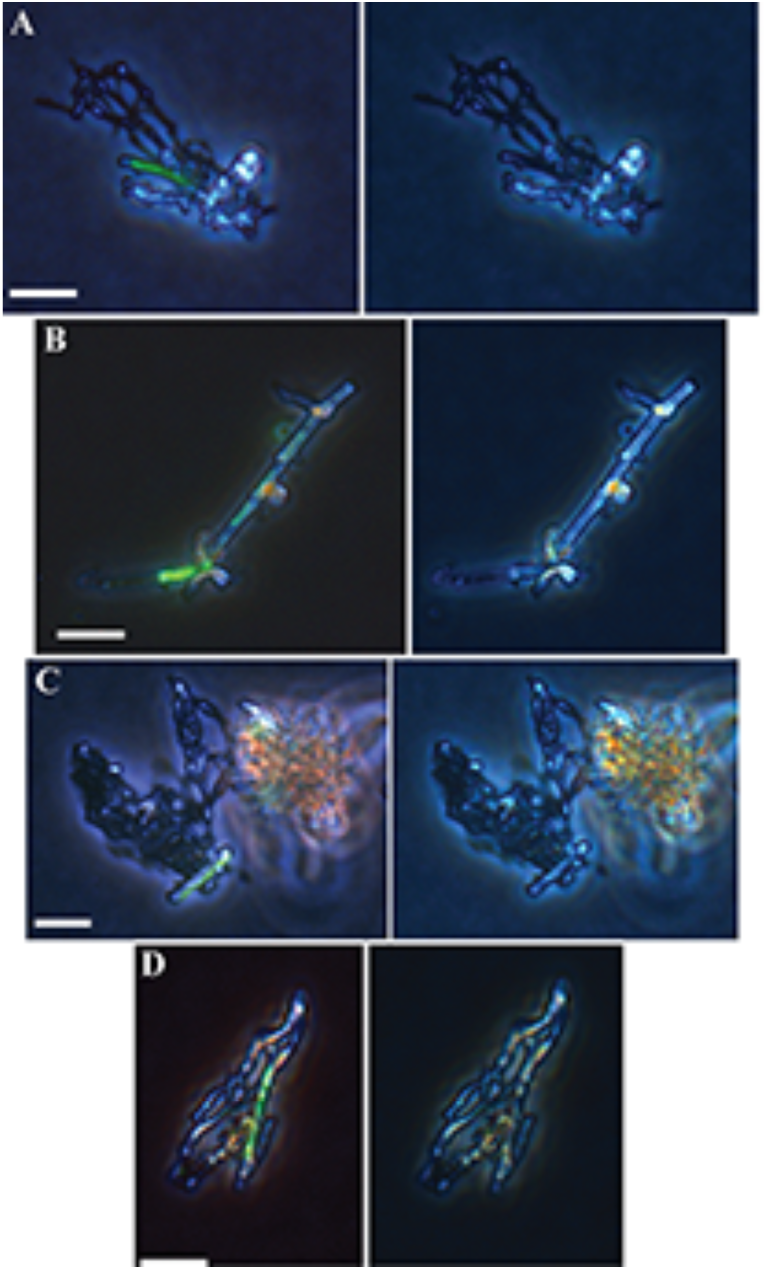
A photo-montage from an in-situ microscope slide of stick-like iron oxides with cells. The left panels are overlays of epifluorescence and phase contrast images showing the juxtaposition of cells and oxides. The right hand panels are the original phase contrast images. The marker bar = 5 μm.

### 3.4 Community Analysis

Amplicon-based analysis of the SSU rRNA gene was used to assess the relative abundances of different bacterial populations within the chambers, as well as assess overall diversity between sites, temporally and spatially, and contrast populations in freshly formed iron mats to bulk iron mat communities. All the chamber samples were dominated by Zetaproteobacteria. The shorter term deployments had relative abundances of Zetaproteobacteria that were between 81 and 87% of the total sequences, while in the two 256h deployments at NHR, the relative abundances were 74 and 31%, Table 3. The communities from three of the chambers (1,2, and 7) from NHR clustered closely together, and grouped with a large cluster of samples from both NHR and S. Hiolo Ridge, while chamber 6 did not cluster with the other NHR samples, (Supplemental Fig. 4). We speculate that the period chamber 6 was not in vent flow may have resulted in colonization of the iron oxides in the cassettes by other, non-FeOB. The microbial communities in the two chambers (5 & 8) at Pohaku clustered with a group of six community iron mat samples, five of which were from Pohaku, indicating that the organisms colonizing the chambers were most closely related to those found in the iron mats at this site (Supplemental Fig. 4). Despite having relatively low abundances of recognizably biogenic oxides, the Pohaku chambers still had high percentages of Zetaproteobacteria, indicating that not all Fe-oxidizing Zetaproteobacteria produce recognizable filamentous morphotypes.

**Table 3.** Relative abundance of Zetaproteobacteria reads compared to total reads from the different chambers based on the SSU 16S rRNA amplicon analysis.

| Chamber | Site | Time(h) | Zeta-Reads | Total<br>Reads | %<br>Zeta-Reads |
| --- | --- | --- | --- | --- | --- |
| 1 | NHR | 148 | 1006 | 1209 | 83.2 |
| 7 | NHR | 148 | 2800 | 3237 | 86.5 |
| 2 | NHR | 256 | 1541 | 1979 | 77.9 |
| 6 | NHR | 256 | 476 | 1427 | 33.4 |
| 5 | Pohaku | 112 | 2278 | 2610 | 87.3 |
| 8 | Pohaku | 112 | 4473 | 5048 | 88.6 |

A more detailed OTU analysis of the Zetaproteobacteria, (Fig. 6), revealed that all the NHR samples had the same representation of Zeta OTUs, although the relative abundances varied. By contrast, in the two Pohaku samples ZetaOtu 1 and ZetaOtu 2 together accounted for over 90% of the Zetaproteobacteria reads. When the populations in the chambers as a whole were compared to microbial mat samples either from Lō’ihi or the Mid-Atlantic Ridge (MAR) (Fig. 6), ZetaOtu 9 was notable for its absence in the chambers. Notable for its presence in the chambers, but absence in mature mats, was ZetaOtu 11. In general, ZetaOtu 11 is rarely found in marine iron mats (McAllister et al., 2011; Scott et al., 2016), despite there being several stalk-forming isolates from this group, including *Mariprofundus ferrooxydans*. The chamber results suggest ZetaOtu 11 may be a signature early colonizer of iron mats, and due to its stalk-forming abilities could play an important role in the genesis of stable mat communities (Chan et al., 2016).

**Fig. 6.**
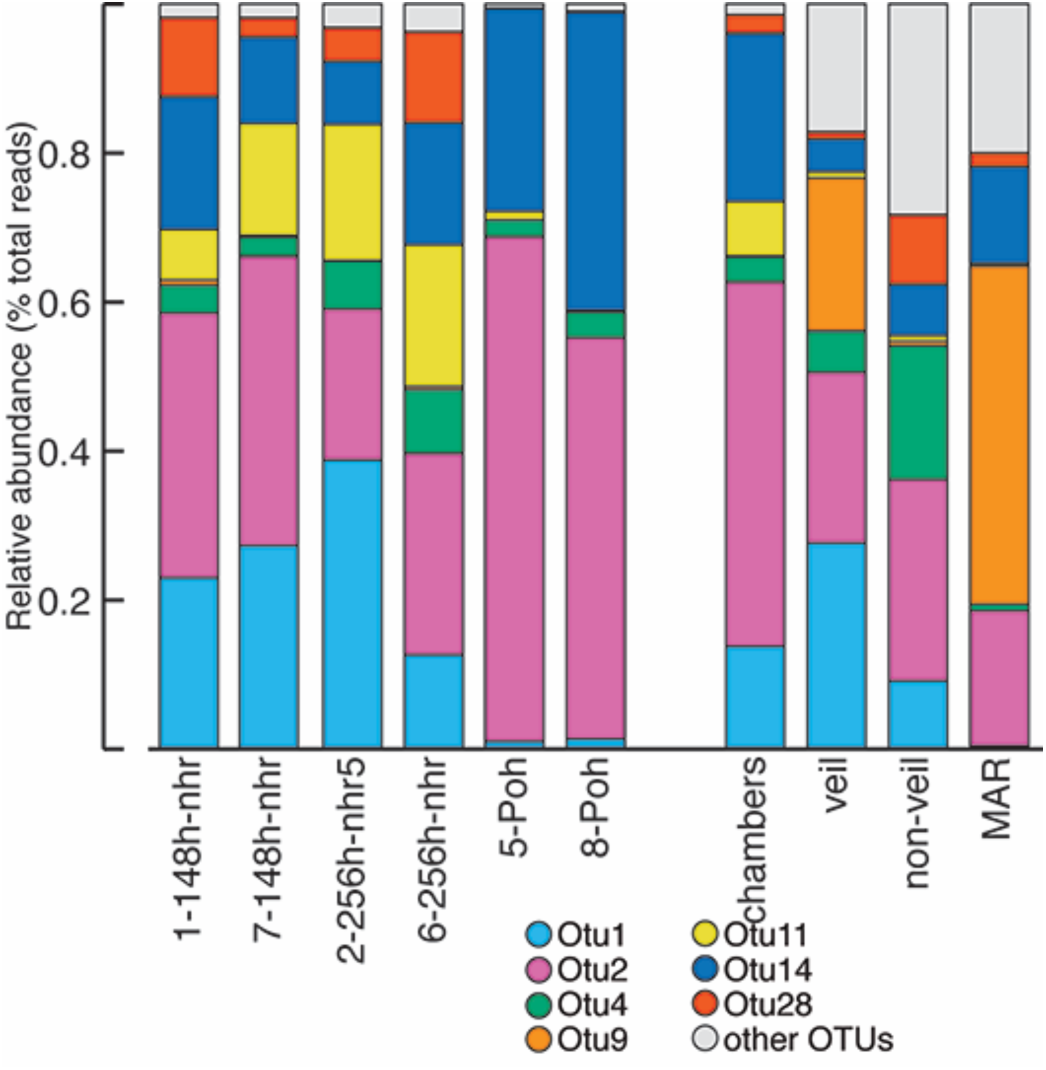
SSU rRNA amplicon community analysis of Zetaproteobacteria present in chambers and iron mats. The left-hand set of columns show the relative abundance of different ZetaOtus from the different incubation chambers. The right-hand columns compare a sample that integrates all the chamber Zetaproteobacteria reads with either veil or non-veil communities in mature Loihi iron mats, or iron mats from the Mid-Atlantic Ridge (MAR). Note the presence of Otu11 in the chambers, and absence from the mats; not the absence of Otu9 in the chambers, but presence in the mats. The data for the iron mat comparisons are taken from Scott et al 2016.

Overall, this community analysis compares and contrasts in interesting ways from an earlier colonization study at Lō’ihi done by Rassa, et al (Rassa et al., 2009). That study evaluated a more diverse group of vent sites at Lō’ihi, both short-term (4 – 10d) and long-term (e.g. 1 year). A different type of colonization chamber that was packed tightly with glass wool was used, and the morphotypes of the FeOB present was not evaluated. Clone libraries (SSU rRNA gene) of short-term incubations were dominated by clones of Zetaproteobacteria from cooler vents (22°C), very similar to what we observed in this study. At the time of the Rassa, et al study, there were also higher temperature vent (64 – 77°C) sites at Lō’ihi, and these had higher abundances of putative sulfur-metabolizing Epsilonproteobacteria (Rassa et al., 2009). Epsilonproteobacteria were only a minor member of the communities we analyzed from these intermediate temperature (40 – 50°) vents that are the hottest fluids currently observed at Lō’ihi. This suggests that as the vents have cooled Zetaproteobacteria have become more prevalent, and is consistent with detectable sulfide being largely absent from the vent fluids in 2013 (Scott et al., 2016.). Terminal restriction fragment length polymorphisms (T_RFLPs) were used in the earlier study to analyze community composition, and an increase in the complexity of the community was noted in long term, versus short term colonization experiments (Rassa et al., 2009). The same trend was noted in this study using amplicon analysis that provides greater resolution of community diversity than T_RFLP; however since only short term deployments were done for this study, it is not possible to make direct comparisons.

### 3.5 Mat accretion

It is important to understand the accretion rates of these iron mats, both for overall microbial mat development, and to determine the potential for biogenic iron mats to serve as an iron source to the surrounding ocean. Recent work from Lō’ihi has shown the primary structural element of the mats are filamentous stalks, or sheaths (Chan et al., 2016). The stalks observed on colonization slides then can serve as a proxy for estimating mat accretion rates of stalk-dominated mats. The most rapid rate of stalk formation observed on a colonization slide was estimated at 2.6 μm/h. This rate is based on the assumption that the cells attached shortly after the slide was deployed and were continuously present until the chamber was retrieved. A stalk production rate of 2.2 μm/h was reported for a pure culture of *M. ferrooxydans* (Chan et al., 2010) based on timelapse imaging, thus the *in situ* estimate is in the same range as the *in vitro* rate. Based on this analysis, if we assume a stalk production rate of 2.5 μm/h, cells that colonize a surface and grow uniformly should accrete at a rate of 60 μm/d, or approximately 420 μm/week. This works out to around 2.2 cm/yr. This is a necessarily simplistic extrapolation of a complex process, i.e. in an actual mat there are multiple colonization events, and it’s unlikely the cells exhibit long-term, uniform growth rates. Nonetheless, it provides an approximation of how fast entire, vent-associated iron mat structures may grow. Interestingly, these estimates for the growth of marine iron mats are substantially slower than the accretion rates of up to 2.2 mm · d^−1^ measured in a freshwater microbial iron mat (Emerson and Revsbech, 1994). The stalk-forming freshwater FeOB *Gallionella ferruginea* can produce stalk at rates up to 80 μm/h, or nearly 40 times faster than *M. ferrooxydans* (Hanert, 1973). Thus, it is reasonable to assume that freshwater iron mats can accrete much more rapidly than marine iron mats.

To further determine if our estimate for the growth rates of marine iron mats is realistic, we did an intentional mat removal experiment at Lō’ihi, by suctioning away a beehive shaped mat that was approximately 30 cm tall and fed by a single diffuse flow orifice on the ocean floor (Supplemental Fig 4). The site was monitored twice more during our expedition, with the last visit coming 10d after removal. While there appeared to be subtle changes at this site, there was not an obvious, visible amount of mat accretion over 10d (Supplemental Fig. 5). Based on our calculations above, the mat would only accumulate to a thickness of <1 mm during these short-term observations. This would be less than could be detected visually with the ROV, thus the fact we did not visibly observe accretion in 10 days is consistent with our estimates. This was intended to be a long-term re-colonization experiment, but, as yet, no subsequent observations have been made of this site. At the Pohaku and NHR vent sites when mat was removed during sample collection with the large suction sampler on ROV *Jason*, we did not detect significant regrowth of the iron mats over the course of an expedition, 10 – 12 days. An example is shown in Supplemental Fig. 6. These same sites had been visited and sampled extensively on a prior expedition to Lō’ihi 18 months earlier; however, during the 2013 expedition there was no obvious sign of the previous disturbance, indicating these 2 – 5 cm thick mats had fully recovered in the intervening time, which would be expected based on our estimates. Thus, our estimates for mat accretion based on observations of rates of stalk-formation seem consistent with visual observations of mat accretion at Lō’ihi.

## 4. Conclusions

The *in-situ* growth chambers proved capable of capturing the growth of biogenic iron oxides, and revealed a diversity of morphotypes associated with FeOB. This work provides an initial estimate for *in-situ* iron oxidation rates for marine Fe-oxidizing bacteria, as well as an estimate for the rate at which iron mats can accrete around diffuse flow hydrothermal vents on the seafloor. An amplicon-based sequence analysis of newly formed Fe-oxides in the chambers revealed Zetaproteobacteria as the dominant phylotype, with a relative abundance as high as 89%. There was a trend towards decreasing relative abundance of Zetaproteobacteria, and increasing overall diversity with incubation time. Colonization slides revealed abundant evidence for colonization by stalk-forming and Y-producing FeOB, as well as a novel stick-like oxide that appeared to be a direct product of microbial iron oxidation. Analysis of the Y-producing microbes using FISH revealed they are members of the Zetaproteobacteria.

## Acknowledgements

We thank the Captain and crew of the RV *Thomas Thompson*, and the ROV *Jason 2* team for their expertise, and help in making this work possible. This work was funded in part by NSF grant OCE-1155754, and NASA Exobiology grant NNX15AM11G.

## Supplemental Figures

Supplemental Fig. 1. An image showing the placement of in-situ incubation chambers (1,2,6, and 7) on diffuse vents associated with NHR.

Supplemental Fig. 2. Temperature graphs from the MTRs placed at NHR and Pohaku. The arrows denote time of deployment and recovery.

Supplemental Fig. 3. A photomosaic from an *in situ* microscope slide of filamentous stalk formation. This represents colonization events by several individual stalk-forming cells. The marker bar = 10 μm.

Supplemental Fig. 4. MED comparison of the total bacterial community composition of the different chambers with 49 iron mat samples collected from different vents at Loihi or from the MAR. The yellow colored panels (top) and arrows (bottom) indicate chambers. The data from other vent sites is taken from Scott *et al* 2016.

Supplemental Fig. 5. Mat removal experiment, a beehive shaped iron mat, left panel, had a black frame placed around it and the mat was removed using the large suction sampler on the ROV *Jason*, and then monitored for 10d. During this time there was little re-growth of the mat. The white bar in the left hand panel = 10 cm, the black frame is approximately 40 cm on each side.

Supplemental Fig. 6. Before and after images from mat sampling on NHR. The left hand panel was taken just prior to sampling with the large suction sampler on ROV *Jason* on 3/19/13, and the right hand image was taken 11d later. A primary vent orifice can be seen as the gray-colored mineral near the bottom of the left hand image. The white circle denotes the approximate sampling area, note there does not appear to be obvious regrowth of the mat during this time.

